# NORMA 3.0: aesthetic and group-aware visualization and analysis of annotated biological networks

**DOI:** 10.64898/2026.11.03.763961

**Authors:** Fotis Baltoumas, Evangelos Karatzas, Mikaela Koutrouli, Alexandros Galaras, Eleni Aplakidou, Iro N. Chasapi, Maria N. Chasapi, Panagiotis Lazanas, Efthimia Lamari, Nataša Pržulj, Ilias Georgakopoulos-Soares, Georgios A. Pavlopoulos

## Abstract

**Summary:** Interpreting a biological network often rests on the relationship between its connectivity and the functional groups annotated on it, such as pathways, complexes, or enriched terms. For example, whether a group forms a cohesive module, how groups overlap, and which proteins connect them. These relationships are often hard to assess visually when groups are numerous or overlap, and most tools represent membership only by node color. NORMA 3.0 is a complete client-side rewrite of its predecessors, delineating and laying out groups with convex hulls, bubble sets, or fog clouds. Multi-group nodes are rendered as pies and directed, and parallel edges retain their evidence channels. Eighteen 2D and nine 3D layouts are scored by silhouette width, outline intrusion and overlap, and benchmarked for group separation and run time. The implementation also includes edge bundling, network importers for six databases, 21 topological measures, group-level and enrichment statistics, comparison of up to ten networks, export of high-resolution images, and an Application Programming Interface.

**Availability and implementation:** NORMA 3.0 is freely available at https://normaviz.org. The source code is available at https://github.com/PavlopoulosLab/NORMA.

**Supplementary information:** Supplementary data are available at *Bioinformatics* online.

## 1 Introduction

Networks fetched from widely known PPI interaction databases (Baltoumas *et al*. 2021) are often combined with functional annotations to test whether proteins in the same pathway, complex, or compartment are also connected. Platforms such as Cytoscape (Shannon *et al*. 2003) and Gephi (Bastian, Heymann, and Jacomy 2009) can show these annotations, but mostly through node color, which fails when there are many groups or when groups overlap (Pavlopoulos *et al*. 2017). NORMA 1.0 was designed to address this problem (Koutrouli *et al*. 2022). It draws each group as a shaded convex hull, turns nodes belonging to several groups into pie charts, and offers igraph layouts.

NORMA-2.0 adds three layout strategies that separate annotated groups (Karatzas *et al*. 2022). The “*virtual node*” strategy adds an invisible hub per group and links it to all members, so the layout draws each group around its own center. “*Group gravity*” reweights the edges, strengthening those inside a group and weakening those between groups, so groups contract and separate. The “*super node*” strategy collapses each group into one node, lays out the resulting network of groups, and then places the members of each group around their super node.

Both versions had clear limits. Every input became a simple undirected graph, edges were straight lines, group membership was shown only by color, and there was no way to measure whether a layout separated the groups. Each user also needed a server-side R/Shiny session. At the same time, previous versions could not serve all cases. For example, STRING scores associations in seven evidence channels (Szklarczyk *et al*. 2025), OmniPath provides signed and directed interactions (Türei *et al*. 2021), and IntAct reports several interaction types for the same protein pair (del Toro *et al*. 2022).

Here we present NORMA 3.0, a complete rewrite that does all its computation in the browser. It keeps the file formats, annotation views, and strategies of its predecessors, but adds new ways to draw groups. It comes with 18 two-dimensional and nine 3D layouts, scores for group separation, support for parallel and directed edges, database importers, group-level analysis, network comparison, a REST API, and exports for other tools (Figure 1 and Supplementary Table S1).

**Figure 1.**
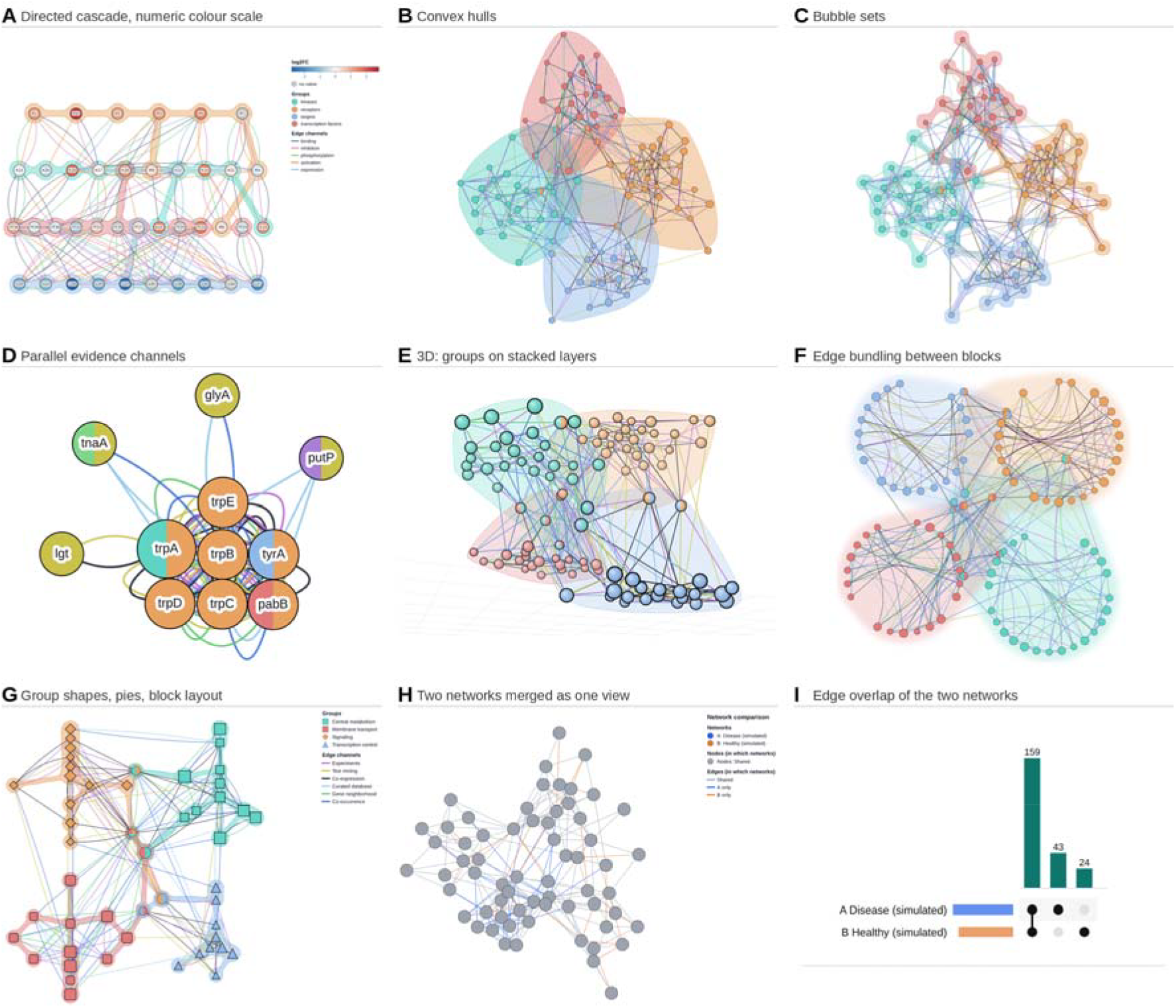
Visual and layout functions of NORMA 3.0. (A) A simulated directed signaling cascade (40 nodes, 155 edges in five channels, four groups) with a hierarchical layout, arrows, fanned Bézier edges coloured by channel, bubble set shading, a diverging log2 fold change scale and a legend. (B, C) Four overlapping modules (100 nodes, 604 edges in six channels) with a weighted force-directed layout. Convex hulls (B) overlap and cover non-members, whereas bubble sets (C) follow each module’s members. Nodes in two modules are two coloured pies. (D) The *E. coli* trp operon (11 genes, 98 edges in six evidence channels) with parallel edges fanned apart in the colors of their channels. (E) The network of B on the 3D Network page, with each group on its own layer. (F) The same network with the groups arranged on a circle, force-directed edge bundling and fog cloud shading. (G) Four modules (50 nodes) arranged on a grid with concentric local layouts, one node shape per module, pies for shared members and bubble sets. (H) Simulated disease and healthy networks over the same 64 nodes, merged into one view, where blue and orange edges appear in only one network. (I) UpSet plot from the Network Comparison page with the shared (159), disease-only (43), and healthy-only (24) edges.

## 2 Implementation and features

### 2.1 Architecture and data model

NORMA 3.0 has been rewritten from scratch, replacing the original R/shiny implementation with a minimal backend implemented in Python and a frontend primarily implemented in JavaScript. It can be served as a single-page static application for basic functionality, or as a full-fledged live server through Docker and Docker Compose, which also includes the REST API. Networks in 2D are drawn with Cytoscape.js (Franz *et al*. 2023), with an optional WebGL renderer for large networks, while the 3D view uses a custom-written WebGL canvas. Uploaded files, views, and sessions never leave the browser. The REST API is served through the Python backend server, enabling connections to third-party services through GET and POST requests. NORMA 3.0 by default supports networks of up to 10,000 nodes, although this limit can be raised when running locally. All operations, including network loading, topological parameter calculations, clustering, layout computations, and rendering, run exclusively in the user’s browser, with long computations running in the background with progress bars. NORMA 3.0 reads files from its previous versions unchanged, with three additions. An optional “*Type*” column turns every edge type into an edge channel, an optional “*Direction*” column marks directed rows, and expression files may carry numeric columns, such as log2 fold changes. The JSON format holds any number of edges per pair, along with their attributes. A view combines the ticked networks with one annotation and one-color source. It keeps its own positions and settings and remembers 60 steps for undo and redo. The central design change is the selection layer. Every layout, statistic, shading, bundling, comparison, and export works on the shown part of a view, which is the nodes of the ticked groups connected by edges of the ticked channels. Hidden nodes keep their positions.

### 2.2 Layouts

Nine layouts work from the connections alone. These are: a force-directed layout (cose), a deterministic weighted Fruchterman-Reingold layout, Kamada-Kawai, stress majorization started from pivot MDS, and circular, concentric, hierarchical, grid, and random layouts. Layouts by groups (see NORMA-2.0 strategies) work in two steps. First, one disc per ticked group is placed without overlap using one of six arrangements: weighted force-directed, cose, circle, grid, hierarchical, and concentric. Then, each group arranges its members with a local layout, and nodes that belong to several groups end up between their groups. In addition, a ‘*spread control’* slider lets nodes repel or attract each other without changing the overall arrangement. After every layout or node movement, NORMA 3.0 reports three scores for the shown part. These are the mean silhouette width (Rousseeuw 1987) of nodes in a single group, the percentage of nodes that fall inside the outline of a foreign group, and the percentage of the outlined area covered more than once. A benchmark runs all 18 layouts on the current view and tabulates their run times and scores, so the user can pick the layout that separates the groups best by numbers rather than by eye.

### 2.3 Aesthetics

Groups can be shaded as convex hulls, fog clouds, or Bubble Sets contours that follow members and bend around nonmembers (Collins, Penn, and Carpendale 2009). Nodes in several ticked groups are visualized as pie charts with up to 16 slices, and each group can take one of 14 different node shapes, so that group membership and expression can be read at the same time. The palettes include the Okabe-Ito, IBM, and Viridis sets, which remain readable for people with color vision deficiency. Node fill can follow group colors, expression colors, or numeric values on diverging or sequential scales. Users can also use several value columns as time steps, and adjust node size by degree, betweenness, closeness, clustering, or a value. Edges can be drawn as straight lines or as Bézier curves. Parallel edges of different channels fan apart so that every channel stays visible. Edges can also be merged, weighted, labeled, given arrows, or bundled with the force-directed edge bundling of Holten and van Wijk (Holten and Van Wijk 2009), which runs in a background worker and survives as editable curves in exported images. A legend of scales, groups, shapes and channels can be shown on screen and in exports (Figure 1).

### 2.4 Importers, analysis and output

Six importers fetch networks and groupings from various repositories. STRING provides one edge per evidence channel and groups its enriched terms. Reactome pathways (Ragueneau *et al*. 2026) group subpathways. OmniPath (Türei *et al*. 2026) provides signed and directed interactions. NDEx networks (Pratt *et al*. 2015) keep their stored layout. IntAct (del Toro *et al*. 2022) splits interaction types into channels, and GO-CAM models and QuickGO annotations (The Gene Ontology Consortium *et al*. 2026) provide causal networks and GO-term groupings. The Network Profiler computes 21 topological statistics defined in igraph (Antonov *et al*. 2026), plus directed-network statistics and community detection using five algorithms. Group analysis reports the density, conductance, and modularity share of every ticked group, and enrichment uses a hypergeometric test with Benjamini-Hochberg correction. Up to ten networks can be compared by node and edge overlap in Venn and UpSet plots as well as by topology and degree agreement and can be merged into one view. Views are exported as PNG, JPEG, WebP, layered SVG or PDF at up to 768 dpi, as GraphML, GEXF, SIF, Cytoscape JSON or NORMA files, as sessions, and to Arena3D^web^ (Karatzas, Baltoumas *et al*. 2021, Kokoli *et al*. 2023) (now Arena3D 3.0 (Karatzas *et al*. 2026)) with each group as a layer. Other applications can call NORMA 3.0 through a REST API endpoint, data links, or postMessage.

## 3 Conclusion

NORMA 3.0 replaces the server-side R/Shiny architecture of its predecessors with a client-side JavaScript implementation and extends the data model from simple undirected graphs to directed multigraphs with typed edge channels (comparison with previous versions in Supplementary Table S1). Bubble Set isocontours, group-specific node shapes, and pie-chart nodes represent overlapping group memberships without relying solely on color, while fanned Bézier curves and force-directed edge bundling preserve individual evidence-specific connections in dense networks. Layout quality is quantified by mean silhouette width, outline intrusion, and outline overlap. In addition, the integrated benchmark across 18 layouts makes layout selection a measured, reproducible step. Partially like in VICTOR application (Karatzas, Gkonta *et al*. 2021), hypergeometric enrichment, group density, conductance and modularity share, and overlap-based comparison of up to ten networks link visual group structure to statistical evidence. These functions apply directly to common analyses such as mapping differentially expressed genes onto a protein interaction network grouped by enriched pathways, inspecting pathway crosstalk and multifunctional proteins at the interfaces of overlapping modules, resolving signaling directionality in OmniPath or GO-CAM networks, and comparing interactomes between disease and control conditions. Client-side execution eliminates installation and data transfer, and standard graph exports together with the REST API integrate NORMA 3.0 into existing analysis pipelines. Limitations remain. The default size limit is 10,000 nodes (can be raised in the configuration when running locally), group-aware layouts separate groups by design rather than by evidence, and topological statistics treat networks as simple graphs, with directionality analyzed separately. Overall, this version significantly improves NORMA’s core functionality and ease of use, making it a useful tool for complex biological network visualization, annotation, and analysis.

## Acknowledgements

Claude Opus 5 (Anthropic) assisted with optimizing parts of the source code and with drafting and editing parts of the manuscript. The authors reviewed and verified all content and take full responsibility for it.

## Data availability

NORMA 3.0 is available at https://normaviz.org and its source code at https://github.com/PavlopoulosLab/NORMA.

## Supplementary information

**Supplementary Table S1.** Features of NORMA 1.0, NORMA 2.0, and NORMA 3.0.

| **Feature** | **NORMA 1.0** | **NORMA-2.0** | **NORMA 3.0** |
|---|---|---|---|
| ***Layouts*** |  |  |  |
| Layouts by connections | 10 (igraph)<sup>a</sup> | 10 (igraph)<sup>a</sup> | 9<sup>p</sup>, incl. weighted FR, Kamada-Kawai, stress majorization |
| Group-aware layouts | Virtual nodes | Strategies 1-3 | Strategies 1-3 on any of 9 algorithms |
| Non-overlapping group arrangements | No | No | 6, each with 10 local layouts |
| Spread and group-size control | NR | NR | Yes |
| Layout quality scores and benchmark | No | No | Silhouette, outline intrusion and overlap; 18 layouts |
| 3D layouts | Plotly 3D hull view | As NORMA 1.0 | 9 interactive 3D layouts |
| ***Aesthetics*** |  |  |  |
| Group shading | Convex hulls | Convex hulls | Convex hulls, bubble sets, fog clouds |
| Pie-chart nodes | Yes | Yes | Yes (up to 16 slices) |
| Node shape per group | 1 | 1 | 14 shapes |
| Color-blind-safe palettes | No | No | Okabe-Ito, IBM, Viridis |
| Node colours | Colour file | Colour file | Color file or numeric values (8 scales), time steps |
| Node size | NR | NR | Data, 6 topological measures or values |
| Labels | NR | NR | Size, color, position, zoom-dependent hiding; edge labels |
| Edge drawing | Straight | Straight | Straight or Bézier curves (fanned parallel edges); merged, weighted, labeled; arrows |
| Edge bundling | No | No | Force-directed (FDEB) in a background worker; up to 10,000 edges |
| Legend (screen and image) | No | No | Yes |
| Themes | Light, dark | NR | 6 |
| ***Data and access*** |  |  |  |
| Parallel edges (channels) | Removed on upload | Removed on upload | Yes |
| Directed and mixed networks | No | No | Yes |
| Database importers | No | No | STRING, Reactome, OmniPath, NDEx, IntAct, GO |
| Programmatic access | No | No | REST API, data links, postMessage |
| Where data are processed | R/Shiny server | R/Shiny server | Browser; optional Python server |
| Size limit | 10 000 nodes | 10 000 nodes | 10 000 nodes |
| ***Exploration and analysis*** |  |  |  |
| Multiple independent views | No | No | Yes |
| Undo and redo | No | No | Yes (60 steps per view) |
| Search, neighborhoods, shortest paths | No | No | Regular expressions; 1-3 steps; all shortest paths |
| Selection-aware functions | Group highlighting | Group highlighting | All functions on ticked groups and channels |
| Topological statistics | 10 | 10 | 21 + directed statistics |
| Group statistics, network of groups, enrichment | No | No | Yes |
| Network comparison | Bar charts of measures | As NORMA 1.0 | 2-10 networks; Venn, UpSet, merged network |
| ***Output*** |  |  |  |
| Image export | Figures<sup>c</sup> | As NORMA 1.0 | PNG, JPEG, WebP, SVG, PDF; up to 768 dpi |
| Interoperable files and sessions | NR | NR | GraphML, GEXF, SIF, Cytoscape JSON; Arena3D<sup>web</sup>; sessions |

<sup>a</sup>Fruchterman-Reingold (FR), Kamada-Kawai, Reingold-Tilford, LGL, Graphopt, GEM, Star, circle, grid and random. <sup>b</sup>Force-directed (cose), weighted FR, Kamada-Kawai, stress majorization, circle, concentric, hierarchical, grid, and random. <sup>c</sup>Export formats not specified in the publication. NR, not reported in the corresponding publication.

